# Chemo-Mechanical Regulation of Tau Phosphorylation Following Traumatic Brain Injuries

**DOI:** 10.1101/2023.07.13.548916

**Authors:** Aayush Kant, Nikhil V. Medhekar, Tanmay K. Bhandakkar

## Abstract

Traumatic brain injuries are characterized by damage to axonal cytoskeletal proteins. Here, we present a mathematical model predicting the chemo-mechanical disruption of intra-axonal microtubule assembly in terms of hyperphosphorylation-led dysfunction of tubulin-binding tau proteins. Intracellular calcium accumulation following a trauma leads to calpain activation, disturbing the downstream kinase-phosphatase activity balance which causes tau hyperphosphorylation. We develop a computational framework, using finite element methods, predicting the spatiotemporal evolution of mechanical stress and ensuing tau hyperphosphorylation in the human brain after traumatic brain injury-inducing loads. We compare our predictions with previously reported experimental and clinical observations to validate the model. Our model provides important insights into the secondary effects of traumatic brain injuries and can be essential in their clinical management.

## 1. Introduction

Traumatic Brain Injuries (TBIs) are a leading cause of injury-related death and disability [1] and more so in low to medium income countries [2]. Over 90% of the incidences of TBI reported clinically present symptoms of a mild TBI (mTBI) or concussion [1], while moderate and severe TBIs especially contribute to the TBI related fatalities [3]. The effects of TBIs are generally observed in two stages, a primary insult which involves an instantaneous mechanical damage, and secondary insults which comprise of a series of neurochemical reactions which occur at a much delayed stage [4]. Commonly mTBIs exhibit a pathological feature called diffuse axonal injury (DAI), the severity of which has been clinically correlated to the extent of disability following the injury [5–8].

DAIs are commonly characterized by damage to white matter axons, which can be observed in-vivo as varicosities or swelling in axonal tracts [8, 9]. Such varicosities are hypothesized to occur due to disruption of axonal transport mechanisms [5, 7]. The axonal cytoskeletal structure comprises a microtubule (MT) core, a bundle of elongated polymers forming a fast transport conduit along the length of axon, surrounded by a periodic actin-spectrin complex and neurofilament proteins all of which collectively contribute towards the axon tensile strength [10–16]. MTs being the stiffest components of the axonal cytoskeleton have been identified to be especially vulnerable to dynamic mechanical rupture with co localized varicosity features observed in-vivo after TBIs [9]. The MTs are assembled into stabilized bundles via cross linking by microtubule-associated protein (MAP) tau proteins [17, 18], which can also undergo a brittle mechanical failure resulting in loss of the MT bundle integrity [11, 12, 18].

Primary injuries, in terms of concomitant mechanical stress and deformation, have been extensively simulated over the brain tissue via finite element (FE) modeling (reviewed in [19–21]). At a scale of individual axons, mathematical models of the axonal ultrastructure have been developed to elucidate the mechanical rupture of axons after stretch injuries. Using a discrete bead-spring model, Peter & Mofrad [18] simulated the collapse of MT bundles due to mechanical failure of cross linking tau proteins using a discrete bead-spring model. Strain-rate dependent viscoelastic contribution tau proteins towards microtubule bundle rupture was modeled by Ahmadzadeh *et al*. [11] and further refined to incorporate the kinetics of tau-tubulin binding [12]. Sendek *et al*. [22] simulated a randomized removal of tau proteins and the resulting MT bundle collapse in a 2D space by equilibriating through steepest descent relaxation. Transfer of mechanical loads due to primary effects of TBI have also been shown to result in microtubule destabilization in three-dimensional axon ultrastructural models [23–26].

However, in conjunction with such primary insults of mechanical nature, neurochemical insults involving a calcium ion (*Ca*^2+^) influx mediated cytoskeletal protein modifications post TBI are also known to debilitate the long-term structural and functional integrity of the axons [5, 10, 27]. Very few computational studies deal with the influence of mechanical stress during the secondary phase of TBI. Kilinc [28] studied the post TBI *Ca*^2+^ evolution due to mechanoporation by explicitly varying the PM permeability. We have previously developed a model utilizing mechanosensitive ion transport channel to predict the intracellular *Ca*^2+^ evolution in presence of a mechanical impulse loading over a two-dimensional (2D) human brain under real-world kinematic loads [29, 30]. Kant *et al*. [13] studied the mechanistic role of spectrin and myelin degradation on increasing the vulnerability of axons towards repeated incidences of mTBIs. Despite widespread evidence of active neurochemical phenomena working in tandem with biomechanical injury observed clinically via TBI biomarkers such as phosphorylated tau, neurofilament residues and spectrin degradation products [31–33] there is no deterministic mathematical link between the occurrence of a TBI and the resulting chemo-mechanical axonal rupture occurring over the time-scales beyond primary mechanical insult.

Here, we propose a new mathematical model which bridges the gap in the literature by shedding light on the mechanistic link between the physiological incidence of TBI and its long term effects. Since tau pathologies have been associated strongly with TBIs, especially moderate and repeated incidences [32, 34], we focus on the hyperphosphorylation-based loss of tau proteins from the MT bundles leading to its destabilization. The model captures the downstream effects of increase in intracellular *Ca*^2+^ concentration due to extra-axonal mechanical load via activation of cysteine protease calpain leading to kinase truncation and lastly imbalance in the tau phosphorylation levels. The model incorporates physiologically relevant parameters which we evaluate by calibrating our model based on previously reported in-vitro and in-vivo experiments [35–37]. Using the model over a 2D brain geometry experiencing in-silico TBI-relevant kinematic loads, we predict the spatial variation in tau hyperphosphorylation in a human brain after TBI. Our model predicts that mechanosensitive imbalance in intraneuronal kinase-phosphatase activity can result in irreversible tau hyperphosphorylation. Our model also predicts that although single mTBI might not cause physiologically relevant tau hyperphosphorylation, repeated mTBIs become progressively more dangerous resulting in unrecoverable MT damage.

## 2. Methods

### Overview of tau phosphorylation pathway

To investigate the extent of secondary injuries after an incidence of a TBI we have developed a mathematical model for predicting the hyperphosphorylation of tau that occurs in response to extra-axonal mechanical pressure loads. Tau protein has multiple (around 30) [36] phosphorylable sites, as shown schematically in Figure 1a. Under homeostatic conditions, some of these sites stay phosphorylated with a dynamic balance of the phosphorylation levels maintained via the activities of kinase (e.g. GSK-3*β*, PKA) and phosphatase (e.g. PP2A) [38] (Figure 1b). Under tauopathic conditions, increased kinase activity results in an increased level of tau phosphorylation (Figure 1b) [38].

**Figure 1:**
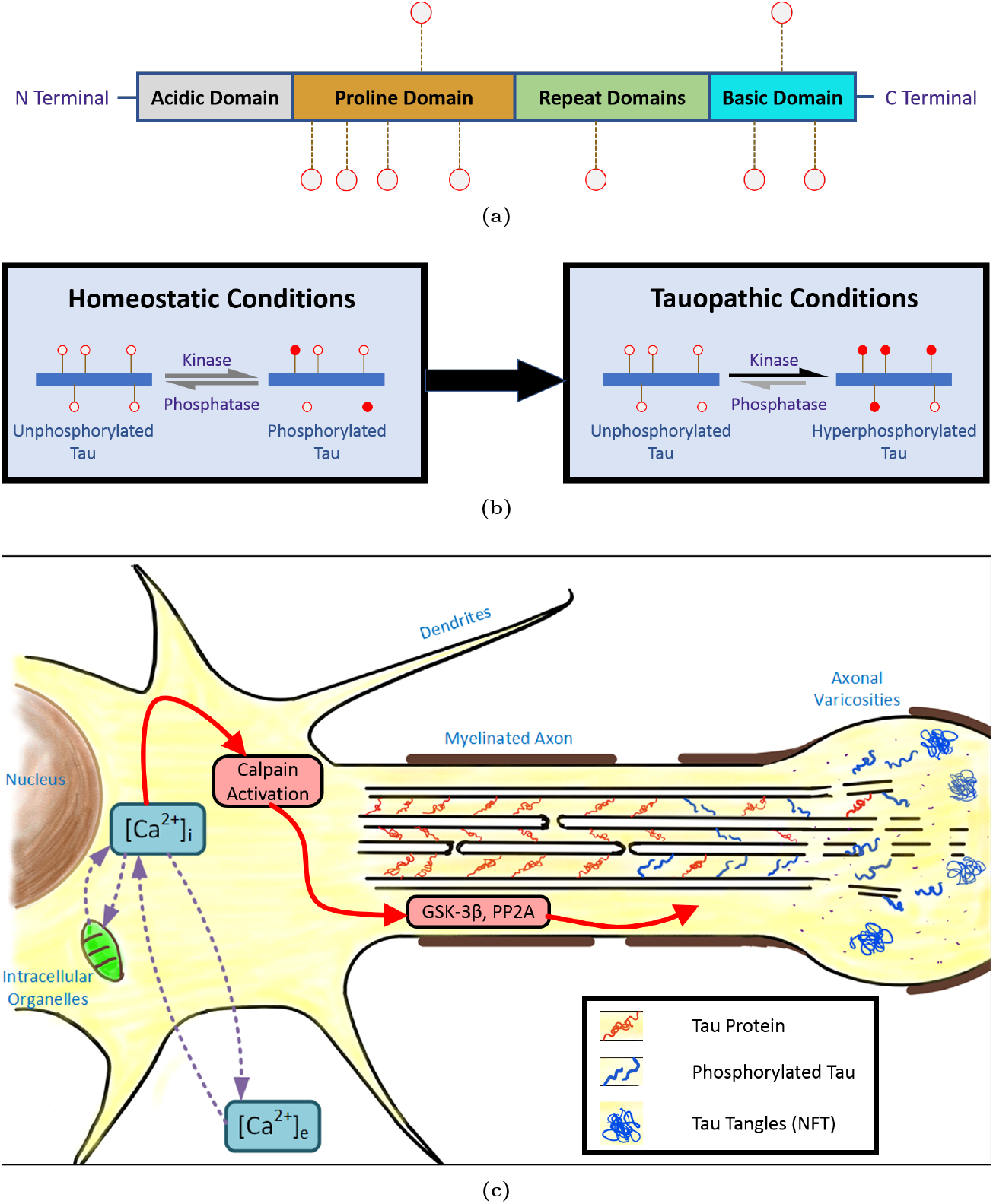
(a) A schematic representation of the structure of tau protein showing multiple phosphorylation sites (b) Under homeostatic conditions, the tau phosphorylation carried out by kinases and dephosphorylation carried out by phosphatases balance each other so as to maintain a constant tau phosphorylation level in the neuron. When a tauopathy occurs, the kinase activity increases and phosphatase activity reduces resulting in hyperphosphorylation of tau proteins. (c) A schematic of pathways leading to rupture of microtubule assembly. *Ca*^2+^ accumulation due to *Ca*^2+^ homeostasis imbalance leads to calpain activation which disturbs the kinase-phosphatase (GSK-3*β* -PP2A) balance resulting in hyperphosphorylation of tau proteins. The hyper-phosphorylated tau proteins thus lose their MT binding abilities, resulting in tau aggregation into neurofibrillary tangles (NFT), and rupture of MT assembly.

Hyperphosphorylated tau are unable to bind to tubulin resulting in the loss of microtubule bundles (Figure 1c). Figure 1c shows a schematic pathway which can result in tau hyperphosphorylation after a TBI. In our model, we incorporate the temporal dynamics of excessive phosphorylation of tau via the disturbed balance between the kinase and phosphatase activity. The imbalance is mediated by the activity of calpain, which breaks down GSK-3*β* into more active fragments. The calpain activity is in turn modulated by changes in intracellular *Ca*^2+^ concentration due to pressure sensitive ion channels in the plasma membrane of the neuron.

### Disturbance in Ca^2+^ *homeostasis*

A compartmental model of *Ca*^2+^ transport between the extracellular space, cytoplasm and the intracellular organelles (for instance the endoplasmic reticulum, ER) can capture the stress induced calcium kinetics in a neuron [29]. We write the rate of change of extracellular *Ca*^2+^ concentration *c*_*e*_ and the intracellular *Ca*^2+^ concentration *c*_*i*_ with respect to time *t* as,

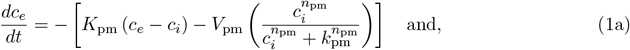

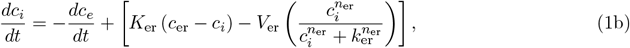

where *c*_er_ is the *Ca*^2+^ concentration inside the ER, and *K*_pm_ and *K*_er_ are the permeabilities of the PM and ER membranes respectively. The last terms in R.H.S. of Eqs. 1a and 1b are a Hill type expression representing the overall *Ca*^2+^ transport via pumps and channels across the PM and ER membranes. *V*_pm_ (*V*_er_) is the maximal overall activity of the PM (ER membrane) pumps and channels, *k*_pm_ (*k*_er_) is the activation concentration for the PM (ER membrane), and the exponent *n*_pm_ (*n*_er_) is the Hill coefficient for the channels in the PM (ER membrane). We previously [29] proposed a mechanosensitive history-dependent rate of ion transport across plasma and ER membranes such that,

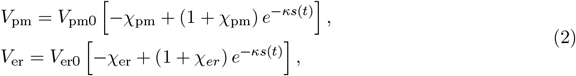

where *χ*_pm_, *χ*_er_ and *κ* are constant parameters, and *V*_pm0_ and *V*_er0_ are the homeostatic values of the coefficients *V*_pm_ and *V*_er_. The stress measure 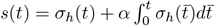 in Eq. 2 comprises of the instantaneous local hydrostatic pressure as well as its time history. Here *α* is a constant parameter representative of the viscosity in the neuron. For a given hydrostatic load *σ*_*h*_(*t*), Eqs. 1-2 can be solved to obtain the evolution of intracellular *Ca*^2+^ concentration *c*_*i*_(*t*). The kinetic parameters appearing in Eqs. 1-2 have been calibrated previously [29] and are listed in Table S1.

### Calpain-I Activation

As discussed in Section 1, excessive *Ca*^2+^ accumulation in the neurons has been implicated in the activation of calpain enzyme. Of the two forms of calpain ubiquitously expressed in a human neuron, calpain-I has a much lower *Ca*^2+^ requirement for its activation as compared to calpain-II, and is hence more severely implicated in the downstream proteolysis of essential proteins during tauopathies [39–41]. Following this observation, we explicitly base our model on activation of calpain-I only.

A calpain molecule comprises multiple subunits each containing multiple sites for *Ca*^2+^ to bind cooperatively [42–45]. Although a multistep process has been hypothesized for calpain activation by *Ca*^2+^, wherein binding of individual *Ca*^2+^ introduce conformational changes assisting further *Ca*^2+^ binding [43, 44, 46, 47], the lack of observation of intermediary species at time scales relevant to secondary injuries following a TBI indicates the multistep calpain activation to be a much faster process. Thus, at the time-scales relevant to TBI, we assume a single step reaction for calpain activation by accumulated *Ca*^2+^, and write the corresponding mass action kinetics as,

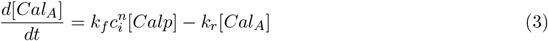

where, *k*_*f*_ and *k*_*r*_ are the forward and reverse rate constants respectively, *n* is the Hill’s constant for the cooperative binding of *Ca*^2+^ ions to a single molecule of calpain, while [*Calp*] and [*Cal*_*A*_] denote the instantaneous concentrations of calpain and activated calpain respectively. In order to conserve the total amount of calpain molecules in the neuron we rewrite Eq. 3 as,

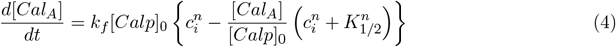

where, [*Calp*]_0_ is the initial calpain concentration, and *K*_1*/*2_ is the *Ca*^2+^ concentration required for half maximal calpain activation, defined in terms of the forward and reverse rate constants as, *K*_1*/*2_ = (*kr/k*_*f*_)^1*/n*^.

The calpain kinetics equation (4) solved simultaneously with the calcium kinetics equations (1-2) allows us to predict the calpain activation following the application of an external hydrostatic stress.

### Calpain Mediated Kinase Truncation

Activated calpain-I can mediate tau hyperphosphorylation via several pathways most implicated of which in-vitro are the increased activities of CDK5 and GSK-3*β* kinases [34]. Although CDK5 activity may increase in response to calpain activation, in vivo evidence for CDK5 being responsible for tau hyperphosphorylation in an Alzheimer’s disease(AD) affected brain is lacking [48–53]. On the other hand, in human TBI, the injury severity is found to correlate with increased activity of GSK-3*β* [54, 55] with ample evidence to implicate calpain in the GSK-3*β* truncation [35, 40, 56–59].

N-terminal truncation of GSK3*β* in-vitro has been reported to be a two-step process with first step generating the larger (40 kDa^1^) fragment *F*_1_, and the second step a smaller (30 kDa) fragment *F*_2_ -both of which are present over a time-scale consistent with secondary insult after a TBI [35, 56]. Accordingly, we model the two step GSK-3*β* truncation process by writing the chemical reactions as,

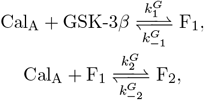

where 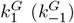 and 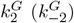 are the forward (reverse) reaction constants for the formation of truncation products *F*_1_ and *F*_2_ respectively. The rate equations for GSK-3*β, F*_1_ and *F*_2_ based on the above chemical reactions are,

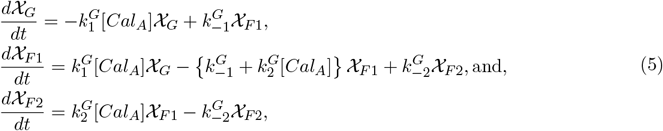

where, *χ*_*G*_ = [GSK-3*β*]*/G*_0_, *χ*_*F* 1_ = [*F*_1_]*/G*_0_ and *χ*_*F* 2_ = [*F*_2_]*/G*_0_ respectively represent the relative volume fractions, measured instantaneously, of full length GSK-3*β*, and its fragments *F*_1_ and *F*_2_ with respect to the initial concentration of GSK-3*β G*_0_. Once the time dependent calpain activation is known, Eq. 5 predicts the time dependent generation of the N-truncated GSK-3*β* fragments (ΔN-GSK-3*β*)

### Tau Phosphorylation/Dephosphorylation

Tau proteins have around 30 phosphorylable sites, of which about 10 sites are known to be hyperphosphorylated in vivo by GSK-3*β* [36, 40, 60, 61]. In post-mortem AD brain neurofibrillary tangle (NFT) aggregates, only 5-9 moles of phosphate per mole of phosphorylated tau (P-tau) are seen [38, 62, 63], suggesting that not all of the available sites may be phosphorylated. In other words, of 2^10^ possible combinations of tau phosphorylation states, around 100 possible states are usually seen in an AD affected brain. This demonstrates the massively combinatorial nature of the phosphorylation/dephosphorylation phenomena which an explicit and deterministic model will most likely be inadequate to capture. Indeed, rule based approaches like agent based modeling have been used in literature to simulate such biological interactions, primarily at the scale of few tau proteins [61].

A single 4 µm long MT bundle in an axon can have 10-100 MTs arranged with a tau protein spacing of ∼ 20 nm [11, 18] suggesting a presence of around 10,000 tau proteins in a single axon. At such a scale, instead of a probabilistic study, we propose a simplified statistically averaged model by making an additional assumption that *the phosphorylation at each distinct site of tau protein follows the same average kinetics*. Although the kinetics for both phosphorylation and dephosphorylation is known to be site-specific, we assert that a numerically averaged kinetics over all the phosphorylation sites can be found [36, 37, 40] at a time-scale relevant to the incidences of secondary insults after TBI. Thus, instead of observing the number of tau proteins being phosphorylated, we account for the status of phosphorylation/dephosphorylation at a site, and compute the average number of sites which are being phosphorylated/dephosphorylated per tau protein. Any unphosphorylated (vacant) site, denoted as ‘ℙ’ can be phosphorylated (occupied), denoted as ‘**P**’ via the full length GSK-3*β*, or the truncation products *F*_1_ and *F*_2_ as,

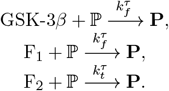

The rate of phosphorylation by full length GSK-3*β* and the intermediate product is assumed to be same due to the lack of sufficient experimental data and to limit the number of parameters in the model. Since kinase truncation increases the phosphorylation effect on tau, 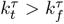. Averaging over the entire axon, we define the total number of phosphorylated sites as N_**P**_ = 𝒩 · [tau]_0_, where 𝒩denotes the average number of occupied sites per tau protein and [tau]_0_ corresponds to the total tau concentration which is conserved over time.

Based on the phosphorylation reactions, the increase in the average number of occupied sites due to tau phosphorylation is,

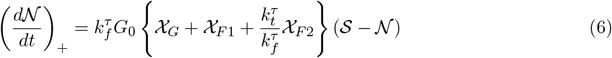

where *G*_0_ is the initial concentration of GSK-3*β* kinase in the system and 𝒮 is the total number of phosphorylable sites on tau. Opposing the phosphorylation, tau dephosphorylation by phosphatases like PP2A have been experimentally observed to follow a Micheles Menten kinetics with a Hill’s coefficient of 1 [37]. Thus, the rate of decrease in average number of occupied sites due to dephosphorylation is,

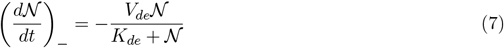

where, *V*_*de*_ is the maximal rate of phosphorylation and *K*_*de*_ is the average site occupancy for half maximal dephosphorylation. Thus, the overall rate of change of the average number of occupied sites per tau protein due to the combined action of phosphorylation by GSK-3*β* and its truncated products (equation 6) as well as dephosphorylation by PP2A (equation 7), is,

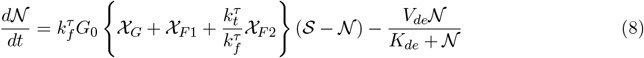

Through the set of equations (1-2, 4, 5, 8) we have built a mathematical framework that allows us to predict the local intracellular *Ca*^2+^ evolution, the resulting calpain activation, GSK-3*β* truncation and tau phosphorylation behaviour, if the evolution of local hydrostatic stress at any point of interest is known. As discussed previously, our model predicts the extent of tau phosphorylation via the parameter 𝒩denoting the average number of phosphorylated sites per tau protein. In the Supplementary Information (Section S1) we present a stochastic interpretation wherein 𝒩 can be used to estimate the number of hyperphosphorylated tau proteins (Equation S2). Although a stochastic prediction, our coarse-grained average interpretation is suitable for predicting the stability of MT assembly in an axon, making our model more viable for clinical applications.

## 3. Results

### 3.1. In-vitro observations allow calibration of model parameters

We have developed a mathematical model to link the physiological occurrence of TBI and the associated primary mechanical loads on the brain tissue with the secondary insult of hyperphosphorylation of tubulin cross-linking tau proteins. We have first linked the extra-axonal mechanical pressure load with imbalance of *Ca*^2+^ in the cytoplasm due to activation of mechanosensitive ion channels and active pumps. Next we quantify and simulate the activation of calpain in the cytoplasm due to intra-axonal *Ca*^2+^ accumulation. Calpain activation results in fragmentation of kinase GSK3-*β*, which increase the kinase mediated phosphorylation of tau proteins. Thus, using our model for a given peri-axonal pressure load we can predict the quantitative extent of tau hyperphosphorylation which is a direct measure of the extent of secondary insult post-TBI. A schematic of the pathway quantitatively simulated is shown in Figure 1c.

Here, we briefly discuss how the parameters associated with each step of the pathway -modeled as a reaction kinetic equation -are calibrated with respect to in-vitro observations reported previously in the literature. The parameters associated with the calcium kinetics in response to mechanical loading, given via Eqs. 1-2 previously [29] calibrated by us are used here and are listed in Table S1 in the Supplementary Information.

Equation 4 governing the calcium mediated calpain activation demands the knowledge of parameters *n, K*_1*/*2_ and *k*_*f*_. While the Hill’s coeffient for *Ca*^2+^ association with calpain varies depending on the binding site, based on the prevously reported values, we choose *n* = 5 [42, 46, 47]. Since calpain autolysis requires the presence of 0.6-2 *µ*M of *Ca*^2+^ [64, 65], we choose *K*_1*/*2_ to be 1.5 *µ*M. Based on Eq. 4, we observe that the theoretically predicted time-scale for calpain activation is 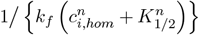, which experimentally is reported to be in the range of 2-10 minutes [65–68].

Thus, we choose *k*_*f*_ = 4.3896 × 10^*−*4^ *µ*M^*−*5^s^*−*1^ to obtain a physiologically relevant calpain activation time-scale of 5 minutes.

Next, we calibrate the kinetic parameters involved in the GSK-3*β* truncation (Eq. 5) against previously reported in-vitro N-terminal truncation of GSK-3*β* by calpain [35]. Under a constant activated calpain concentration, we plot the evolution of the volume fractions of the full length GSK-3*β* and both truncation products in Figure 2a. We observe an excellent qualitative similarity between our predictions and the evolution reported experimentally [35]. The in-vitro GSK-3*β* fragmentation was reported at calpain concentration of 0.2 units/ml (∼13.4 nM) [35]. Setting calpain concentration as 13.4 nM we choose the reaction parameters so as to obtain a good fit for the numerically predicted time-evolution of the volume fractions of full length GSK-3*β* and its first truncation product -fragment F1 (shown in Figure 2a). The numerically predicted volume fraction of the final truncation product -fragment F2 matches the experimentally obtained evolution very closely thereby validating the calibration (blue solid line in Figure 2a).

**Figure 2:**
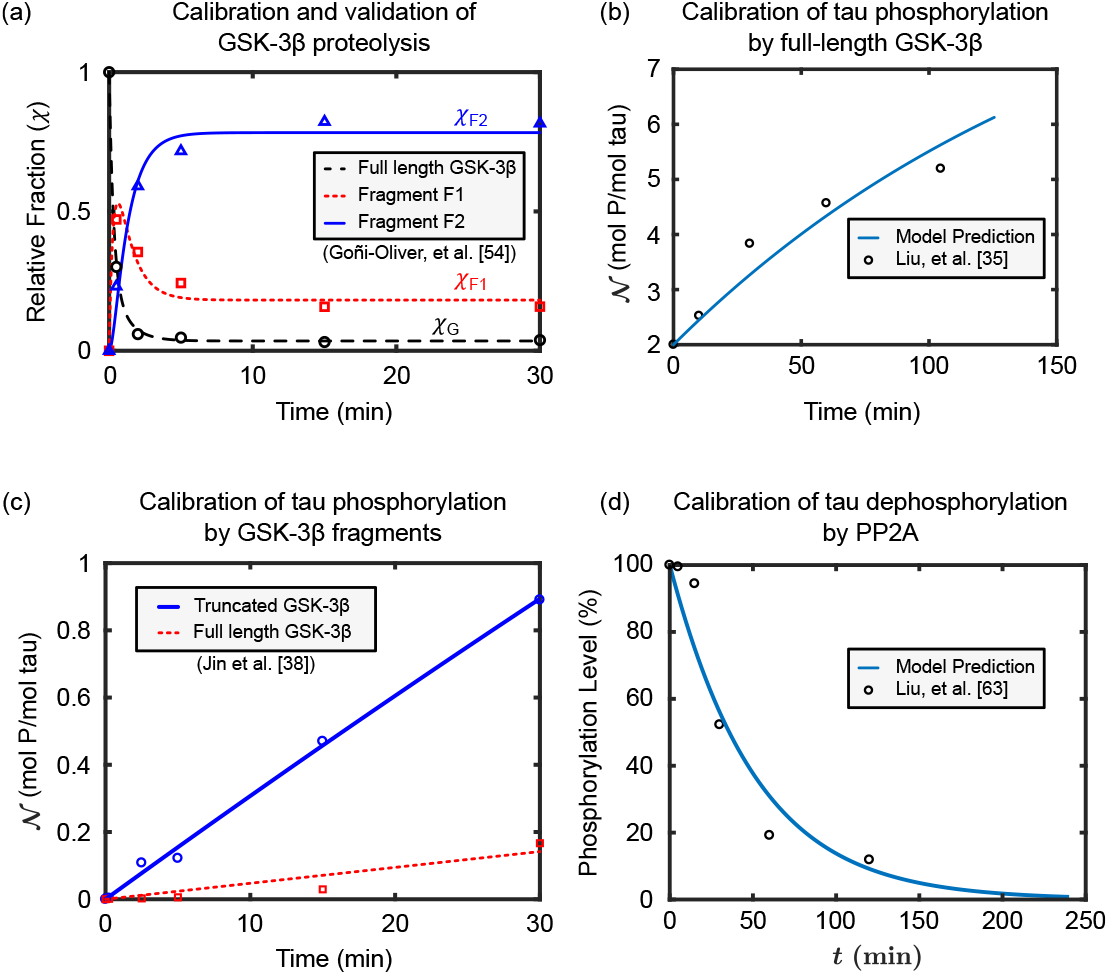
The parameters involved in Eqs. (1-2, 4, 5, 8) are estimated by calibrating the model to experiments reported in literature. (a) Comparison of the evolution of relative fractions of GSK-3*β* and its truncation products *F*_1_ and *F*_2_ using the calibrated values of kinetic parameters (noted in Table 1) in Eq. 5 with the experimental observations of GoÑi-Oliver *et al*. [35] denoted by markers. (b) Comparison of the evolution of phosphorylation of PKA prephosphorylated tau by full length GSK-3*β* predicted by Eq. 6 using the calibrated value of 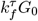 (noted in Table 1) with the values reported by Liu *et al*. [36]. (c) Comparison of the extent of phosphorylation by full length and truncated GSK-3*β* predicted via Eq. 6 using the calibrated value 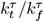 (noted in Table 1) with the experimental values reported by Jin *et al*. [40]. (d) Comparison of the temporal decay of tau dephosphorylation by PP2A computed through Eq. 7 using the calibrated values of parameters *V*_*de*_, *K*_*de*_ (noted in Table 1) with the corresponding experimental result of Liu *et al*. [37].

Further, we calibrate the kinetic parameters involved in tau phosphorylation and dephosphorylation (Eqs. 6, 7). The kinetic parameter 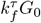 in our model replicates the in vitro increase in susceptibility of tau to GSK mediated phosphorylation in the presence of the kinase PKA [69, 70] in absence of any GSK truncation (set *χ*_*F* 1_ = *χ*_*F* 2_ = 0). The in-silico predictions of tau phosphorylation levels qualitatively follow the experimental observations by Liu *et al*. [36] (Figure 2b). Upon a quantitative comparison with the tau phosphorylation observed by Liu *et al*. [36], as shown in Figure 2b, we choose 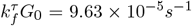. In order to obtain the role of GSK truncation, and calibrate the parameter 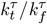 in Eq. 6, we simulate in-silico the experiment by Jin *et al*. [40]. First GSK-3*β* is proteolyzed with 2 nM activated calpain in-silico (Eq. 5) for 10 minutes as done in-vitro. Thus obtained mixture of GSK-3*β* and its fragmentation products is used to obtain the evolution of tau phosphorylation (solid lines in Figure 2c). The in-silico prediction is compared with the kinetics of phosphorylation of S-199 site of tau reported by Jin *et al*. [40] (points marked in Figure 2c) to obtain the value of 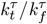.

To validate the evaluated parameters associated with Eq. 6, we numerically replicate the observations made independently by Jin *et al*. [40] where the GSK-3*β* truncation by 1nM calpain-I is allowed to proceed for different times and the subsequent kinase activity is recorded. The same study is carried in silico using Eqs. 5 and 6 of our model with calibrated parameters to predict the time variation of relative kinase activity between the full length kinase, and its truncation products. The result predicted by model and plotted in Figure S1a is qualitatively similar to the result of Jin *et al*. [40] (reproduced in Figure S1b) and succeeds in capturing the increase in the kinase activity post truncation. However the experiments predict that the relative activity doubles in ∼ 30 min [40] while our model predicts that it will triple in ∼ 20 min. The quantitative dissimilarity can be attributed to the fact that the model calibration made use of the work of GoÑi-Oliver *et al*. [35] based on N-truncation of GSK-3*β* while the work of Jin *et al*. [40] which forms the basis of Fig. 2c is based on C-terminal truncation of GSK-3*β*.

Lastly, the dephosphorylation kinetics parameters *V*_*de*_ and *K*_*de*_ are evaluated via a quantitative comparison with the in-vitro PP2A mediated dephosphorylation of site ser-199 on tau protein [37]. The comparison is shown in Figure 2d. The parameter values calibrated and use hereafter are listed in Table 1.

**Table 1:** Parameter Values for Enzyme Kinetic Reactions.

|  | Parameter | Definition | Value |
| --- | --- | --- | --- |
| Calpain<br>Activation | $K_{1/2}$ | $c_i$ for half maximal activation | $1.5\mu\text{M}$ |
| | $n$ | Hill Coefficient | 5 |
| | $k_f$ | Forward Rate Constant for activation | $4.3896 \times 10^{-4}\mu\text{M}^{-5}\text{s}^{-1}$ |
| | $[\text{Calp}]_0$ | Homeostatic Calpain concentration | $0.1\mu\text{M}$ |
| GSK-3β<br>Truncation | $k_1^G$ | Forward rate constant for truncation step 1 | $3.312\mu\text{M}^{-1}\text{s}^{-1}$ |
| | $k_{-1}^G$ | Reverse rate constant for truncation step 1 | $0.0086\text{s}^{-1}$ |
| | $k_2^G$ | Forward rate constant for truncation step 2 | $1.121\mu\text{M}^{-1}\text{s}^{-1}$ |
| | $k_{-2}^G$ | Reverse rate constant for truncation step 2 | $0.0035\text{s}^{-1}$ |
| Tau<br>Phosphorylation/<br>Dephosphorylation | $k_f^\tau$ | Reaction Rate for Tau Phosphorylation<br>by full length $[\text{GSK-3}\beta]_0$ | $k_f^\tau G_0$<br>$= 9.6282 \times 10^{-5}\text{s}^{-1}$ |
| | $G_0$ | Homeostatic concentration of GSK-3β | |
| | $k_t^\tau$ | Reaction Rate for Tau Phosphorylation<br>by truncated product $F_2$ | $k_t^\tau/k_f^\tau = 29.292$ |
| | $V_{de}$ | Maximal tau site dephosphorylation rate | $3.2615 \times 10^{-3}\text{s}^{-1}$ |
| | $K_m$ | Total tau concentration for half maximal<br>dephosphorylation | $11.6\mu\text{M}$ |
| | $[\text{tau}]_0$ | Total tau concentration in neuron | $4.36\mu\text{M}$ |
| | $\mathcal{S}$ | Number of sites on each tau protein | 10 |

### 3.2. Homeostasis predicted in-silico is quantitatively consistent with in-vivo observations

Before making predictions on the extent of tau hyperphosphorylation due to mechanical stress, we first verify that under stress-free homeostatic conditions the kinetics predicted by the model are stable and consistent with what is expected in-vivo.

We have previously [29] established that in absence of any external mechanical stress, the calcium kinetics model (Eq (1-2)) is inherently stable, and there is no net *Ca*^2+^ transport across either the PM or ER membranes. Therefore no *Ca*^2+^ accumulation occurs in absence of any external mechanical stress, and homeostasis is sustained such that the intracellular *Ca*^2+^ concentration is maintained at its initial sub-micromolar levels.

The chosen calpain kinetics parameters permit a negligible calpain activation unless *Ca*^2+^ accumulation occurs. Using the calibrated parameter values listed in Table 1 and Table S1, Eq. 4 yields a negligible fraction of calpain activation under homeostatic conditions 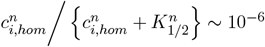 which concurs with the experimentally observed lack of activated calpain under an undisturbed homeostatic condition [45, 65]. Further downstream, following a negligible calpain activation, equation (5) and the associated GSK-3*β* truncation kinetics parameters listed in Table 1 ensure that a negligible fraction (∼ 10^*−*9^) of GSK-3*β* is truncated at homeostasis, allowing it to remain as a full length kinase.

Thus, under homeostasis, in the absence of any mechanical loads, the model predicts a lack of calpain activation and GSK-3*β* truncation. However, even full length GSK-3*β* can cause phosphorylation of tau. Thus, we can expect that even at homeostasis, some level of phosphorylation does occur on the tau proteins. To measure the steady-state homeostatic tau phosphorylation levels, we set *d*𝒩 */dt* = 0 in Eq. 8, thereby obtaining the quadratic equation,

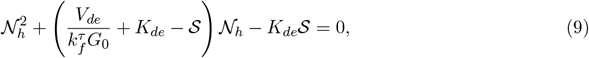

solving which we predict that the average number of occupied sites per mole of total tau at homeostasis 𝒩_*h*_ = 2.33.

This prediction is indeed consistent with in-vivo expectations in homeostatic brain. Under homeostatic conditions, axons do exhibit some phosphorylation of tau in order to promote MT assembly [38]. Post mortem analysis on a normal human brain reveals an average phosphorylation levels of 2-3 moles of phosphate per mole of tau [36, 38, 60, 63]. Thus the homeostatic phosphorylation level predicted by our model 𝒩_*h*_ = 2.33 is well within the in-vivo homeostatic tau phosphorylation levels. The reasonable value of 𝒩_*h*_, obtained with no additional parameter-fitting, further validates the calibration procedure delineated in Section 3.1 and justifies the appropriateness of parameter values used in our model.

In absence of any external mechanical loads, the model predictions are summarized as follows,

- homeostatic intracellular *Ca*^2+^ concentration is maintained,
- negligible calpain activation occurs,
- GSK-3*β* exists in its full length form without getting truncated.
- The tau phosphorylation is maintained at a homeostatic level of 𝒩_*h*_ = 2.33 molP/mol total tau.

In the next section we apply an external load to predict the departure of neuro-chemical response of an axon from the homeostatic condition due to a TBI.

### 3.3. Model predicts increased tau phosphorylation on individual axon after repeated pressure impulse

Having calibrated and validated the model, we next apply realistic pressure impulses to simulate the downstream secondary mechano-chemical effects of a TBI on individual axon, without considering the spatial effects of load transfer in the tissue. For the purpose of in-vitro experiments on brain tissue samples, realistic stress/strain impulses in brain tissue are often idealized in the form of single or repeated rectangular pressure impulses [71–73]. Consequently as a first step, we consider an idealized mechanical load comprising of 6 repeated rectangular pressure impulses, with each impulse of magnitude 10 kPa, duration 10 ms, and a resting period of 50 ms between each successive impulse, as shown in the inset of Fig. 3 to be the input hydrostatic stress *σ*_*h*_ for non-spatial version of our model. We numerically solved equations (1-2, 4, 5, 8) for temporal changes in *Ca*^2+^ concentration, calpain activation, GSK-3*β* truncation and tau phosphorylation/dephosphorylation adopting the parameters listed in Table 1 and Table S1. The resultant intracellular *Ca*^2+^ evolution and the time dependent activation of calpain-I are shown using solid and dotted lines respectively in Fig. 3a. The intracellular *Ca*^2+^ concentration is seen to increase to approximately 14 times the homeostatic value immediately after the action of the pressure impulses. The increased *Ca*^2+^ results in a calpain activation albeit with a small delay, permitting a peak activated calpain fraction of 0.086 ([*Cal*_*A*_]_*peak*_ = 8.6 × 10^*−*3^*µ*M) approximately 2 minutes after the application of mechanical load as shown in Fig. 3a. The increased calpain activation is sustained up to ∼ 30 min after the load application.

**Figure 3:**
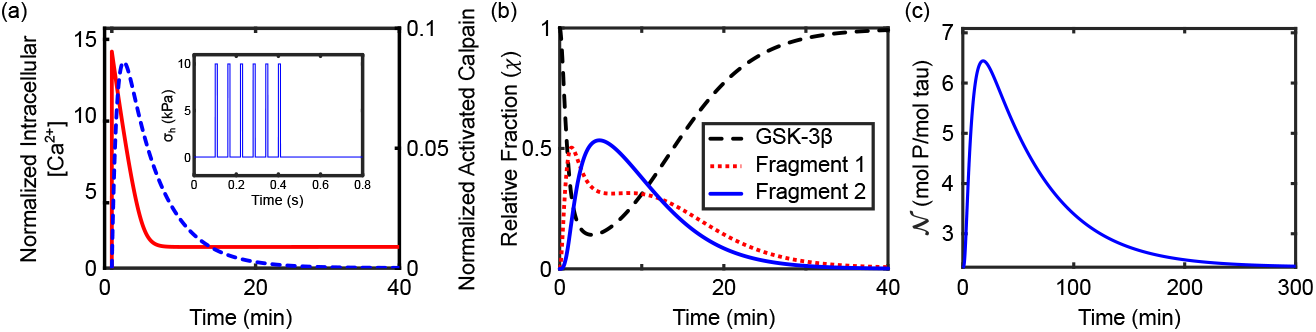
The time evolution of (a) intracellular calcium ion concentration and concentration of activated calpain (b) GSK-3*β* kinase truncation by activated calpain, and (c) tau phosphorylation levels, upon application of 5 rectangular impulses each of magnitude 10 kPa, and duration of 10 ms, as shown in the inset of (a).

The time-scale of calpain activation of ∼ 20 minutes is in close agreement with the experimental observations of notable increase in the activation of calpain-I and formation of calpain specific proteolysis products within 15 minutes [67, 74] and 30 minutes [67, 75] of injury respectively. Further the peak activated calpain concentration [*Cal*_*A*_]_*peak*_ ∼ 8.6 × 10^*−*3^*µ*M predicted in-silico after the representative TBI also falls within the range 1-20 nM generally adopted as calpain concentration in the experiments involving in vivo calpain activity [35, 40, 76]. These observations further serve to validate the choice of parameters listed in Table 1.

Activated calpain in turn leads to truncation of GSK-3*β* kinase progressively into fragments *F*_1_ and *F*_2_. Figure 3b shows the evolution of relative fractions of full length kinase and its products *F*_1_ and *F*_2_ with time. In early stage, the fraction of truncated products increases with time due to ready availability of activated calpain. However once the activated calpain returns to negligible level, the formation of truncation products is no longer be feasible and the full length kinase stays untruncated. It can be noted that the GSK-3*β* truncation follows a similar time scale as calpain activation with GSK-3*β* fragments being formed up to 30 minutes after the stress load. This is expected due to the similar time scales in the experimental observations of GoÑi-Oliver *et al*. [35] which is the basis for the calibration of parameters in the GSK-3*β* truncation equation (5).

The fragmented kinase made available after GSK-3*β* truncation interacts with tau proteins resulting in a higher level of phosphorylation as shown in Fig. 3c. It can be seen that the average number of occupied sites per mole of total tau (𝒩) increases from its homeostatic value (𝒩_*h*_ = 2.33) to a peak value 𝒩_*peak*_. With the progression of time, as the availability of more active truncation products decreases, the dephosphorylation effects of PP2A phosphatase dominates and reduces 𝒩 back to 𝒩_*h*_. This restoration occurs slowly and higher levels of phosphorylated tau sites are seen till ∼ 200 min (∼ 3.5 hours) after the applied pressure load. Thus we note that the time scale of tau phosphorylation/dephosphorylation is much higher as compared to the time scales of calpain activation and kinase truncation. Such high time-scales of elevated tau phosphorylation levels can also be ascertained in the in-vitro experimental observations of Liu *et al*. [36, 37] and Jin *et al*. [40].

### 3.4. Increased tau phosphorylation is predicted in human brain after physiologically relevant loads

In order to invoke spatial dependence in the present work, analogous to our earlier work [30], we next apply the non-spatial neurochemical kinetics model pointwise throughout the continuum of brain tissue. Depending upon the spatio-temporal variation of mechanical pressure, the *Ca*^2+^, calpain, GSK-3*β* truncation and tau phosphorylation at every point in the brain tissue would be governed by the model Eq. (1-2, 4, 5, 8). Such spatial model allows us to assess the effect of non-homogeneous dynamic stress distribution while accounting for realistic loading, tissue geometry and material response on the spatio-temporal prediction secondary damage via tau hyperphosphorylation. To test our present model, we use the experiments by Nusholtz *et al*. [77], who recorded the realistic kinematic response of human brain to TBIs. We have simulated the experiment by creating a 2D structural Finite Element Model (FEM) of a human brain with linear viscoelastic properties, on which we applied the kinematic load in the form of translation and rotational acceleration/deceleration (See [30] for more details). In Table 2 we have listed the translational and rotational acceleration and velocities, and their time durations as observed by Nusholtz *et al*. [77] for three distinct kinematic loading conditions. Using FEM model developed previously [30] in COMSOL Multiphysics for each loading case, we obtain the spatio-temporal variation of hydrostatic stress throughout the brain geometry [30].

**Table 2:** Parameters for the kinematic impulse loading based on the experiment of [77] applied on the two-dimensional brain model [30].

| Case No. | Peak Linear Velocity (m/s) | Peak Angular Velocity (rad/s) | Peak Linear Acceleration (m/s<sup>2</sup>) | Peak Angular Acceleration (rad/s<sup>2</sup>) | Impulse time (ms) |
| --- | --- | --- | --- | --- | --- |
| 1 | 7 | 28 | 1900 | 7250 | 10 |
| 2 | 4.5 | 30 | 420 | 3900 | 25 |
| 3 | 3.8 | 30 | 1350 | 7500 | 12 |

For each of the three load cases described in Table 2 the spatial distribution of the pressure, intracellular *Ca*^2+^ concentration and the extent of tau phosphorylation when each of them reaches their peak values are plotted in Figure 4. We see that for the load case 1, since the peak linear velocity and acceleration are highest, the pressure reaches much higher values than for the cases 2 and

**Figure 4:**
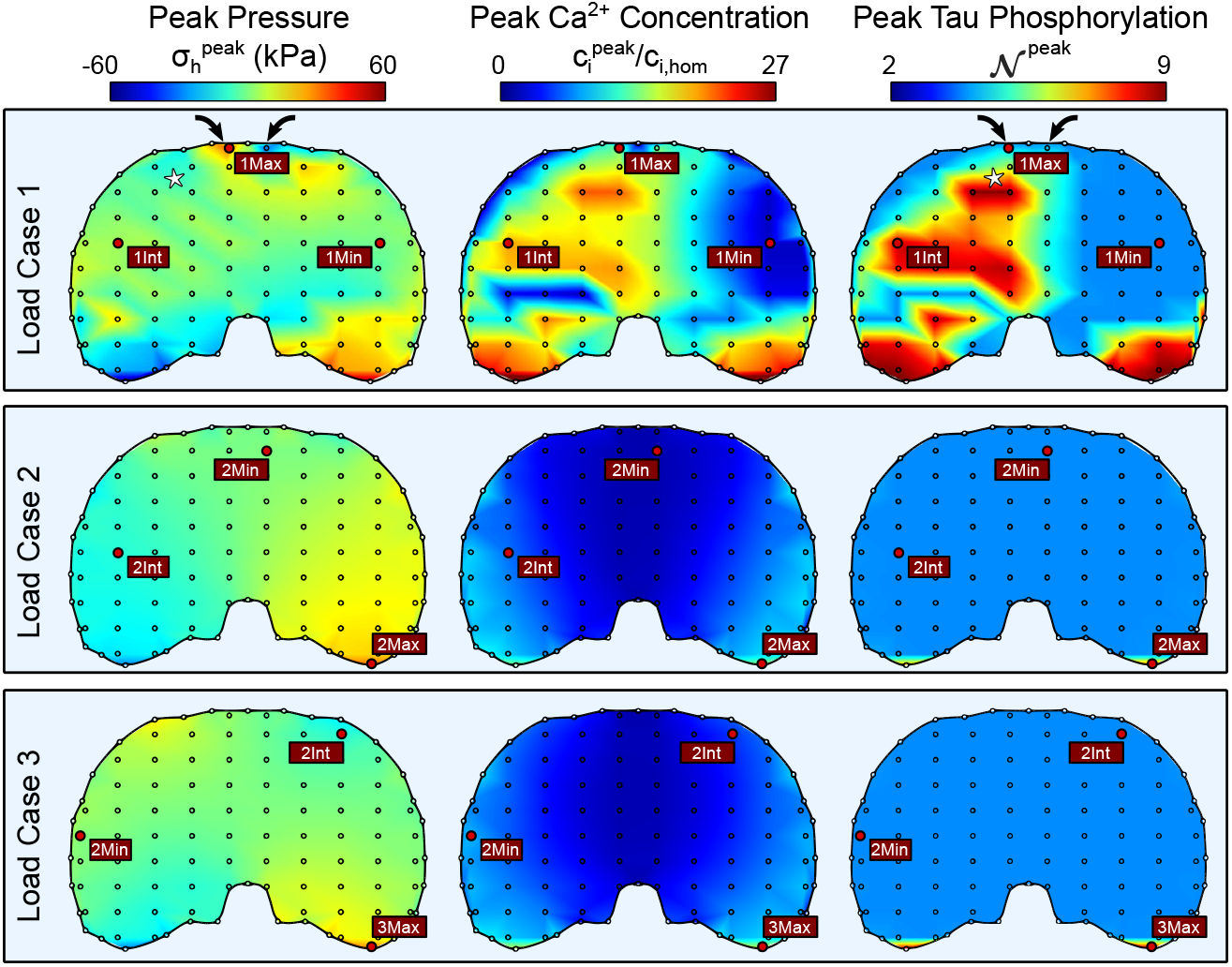
Numerical simulations predict the spatial distribution of pressure at time *t* = 0.02*s* when it peaks, Intracellular *Ca*^2+^ concentration at time *t* = 6*s* when it peaks, and the extent of tau phosphorylation at *t* = 20 minutes when it peaks. The simulations are carried out for the three load cases described in Table 2.

Consequently, the intracellular *Ca*^2+^ concentration as well as the extent of tau phosphorylation are much higher and much more widespread for this case. Indeed the observations by Nusholtz *et al*. [77] suggest that this load results in subarachnoidal hematoma along with frontal lobe hemorrhage implying a moderate to severe TBI. For the other two cases, we predict that the tau phosphorylation levels are much closer to homeostatic levels, rising only near the geometry extremities. Our prediction agrees with the observation of Nusholtz *et al*. [77] which state that case 2 results in no hematoma while case 3 results in no injury. For these cases, the highest value of N_*peak*_ attained is less than observed in vivo phosphorylation levels post fatal tauopathies [38, 62]. These loading cases might correspond to mild TBI, where indeed the tau phosphorylation levels are not expected to reach high values, unless such repeated injuries result in chromatin traumatic encephalopathy (CTE) [8, 78].

Lastly, we observe for case 1 that the points of peak pressure (marked by black arrows, Figure 4) do not necessarily colocalize with the points of peak intra-axonal tau phosphorylation (marked by white asterisk, Figure 4). That is because, as reported by us previously [29, 30], severity of secondary injuries depends not only the peak pressure values but also their durations. As the stress is propagated through the brain tissue, the wavelength of the stress waves, as well as their repeated propagation could play a more significant role than their amplitudes.

In each of the loading case, we identify three locations of interest where the peak stress is maximum, minimum and intermediate and denoted as Max, Min and Int respectively (labeled and marked by circles in Figure 4). For each of the nine stress evolution cases, we use our current model to predict the extent of calpain activation and tau phosphorylation. Table 3 lists the peak hydrostatic stress, peak intra-axonal calcium concentration and the peak value of N (N_*peak*_) for each case at the three locations.

**Table 3:** Local hydrostatic stress, local intra-axonal calcium accumulation and subsequent tau phosphorylation at the points experiencing maximum, minimum and intermediate hydrostatic stress corresponding to the three loading cases shown in Table 2. The points of interest are identified based on the finite element analysis described in our previous work [30].

| Case No. | Point of interest | $\sigma_h^{peak}$ (kPa) | $c_i^{peak}/c_{i,hom}$ | $\mathcal{N}^{peak}$ (mol P/mol tau) |
| --- | --- | --- | --- | --- |
| 1 | 1Max | 32.56 | 13.54 | 3.32 |
|  | 1Int | 3.94 | 19.23 | 8.33 |
|  | 1Min | 1.66 | 1.78 | 2.33 |
| 2 | 2Max | 33.30 | 14.55 | 8.11 |
|  | 2Int | 10.87 | 6.31 | 2.33 |
|  | 2Min | 0.58 | 1.23 | 2.33 |
| 3 | 3Max | 29.99 | 15.91 | 8.63 |
|  | 3Int | 10.66 | 4.53 | 2.33 |
|  | 3Min | 0.16 | 8.08 | 2.33 |

We see that typically the peak levels of tau phosphorylation are higher if the peak *Ca*^2+^ concentration 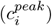 reaches a higher value. However, the relationship between 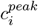 and 𝒩_*peak*_ is highly non-linear. For 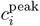 comparable to homeostatic value *c*_*i*,hom_, the peak tau phosphorylation levels 𝒩_*peak*_ does not deviate significantly from the homeostatic value 𝒩_*h*_ = 2.33 as seen from the ‘Min’ location in each of the case. On the other hand at locations away from homeostasis e.g. ‘Max’ location in each case, the tau phopshorylation levels see major increase as compared to 𝒩_*h*_. Table 3 indicates that an intracellular *Ca*^2+^ accumulation even up to ∼8 times the homeostatic *Ca*^2+^ levels, does not increase the tau phosphorylation level much over their homeostatic value (𝒩 _*h*_ = 2.3345). However once the intracellular *Ca*^2+^ reaches ~ 10 times the homeostatic levels, the increase in tau phosphorylation levels are appreciable, and continue to rise faster as the *Ca*^2+^ accumulation increases. The variable sensitivity exhibited by _*h*_ to the extent of deviation of *Ca*^2+^ concentration from homeostatic value provides an interesting insight into the harmfulness of accumulated intracellular *Ca*^2+^ i.e. a limited increase in the *Ca*^2+^ concentration should not be threatening as it may not directly result in an increased tau phosphorylation.

### 3.5. Altered PP2A dephosphorylation results in irrecoverable tau phosphorylation

From Fig. 3 we observe that although the tau phosphorylation levels are elevated after the incidence of mechanical loading, eventually they return to the homeostatic state. The recovery of tau phosphorylation to homeostatis occurs due to tau dephosphorylation by phosphatase PP2A (equation (7)). Thus if the PP2A activity is altered via sensitivity to either primary or secondary insults after TBI, we can expect the recovery to be incomplete such that the elevated phosphorylation levels do not return to homeostatic levels.

In reality, a reduced PP2A activity due to increased presence of phosphatase inhibitors has been reported in tauopathic conditions such as post AD human brain [38] and brains of patients afflicted with severe TBI [54]. Owing to lack of precise knowledge of mechanisms leading to the reduction of PP2A activity after a TBI prevents us from including its effects in our mathematical model. However the consequence of reduced PP2A activity can be phenomenologically simulated by artificially reducing the kinetic parameter *V*_*de*_ in our model (Eq 7) which signifies the maximal activity of the phosphatases towards phosphorylated tau. As a representative example, for this simulation we chose the loading case 2 listed in Table 2, and further select the point where the peak stresses are in an intermediate range (CASE 2Int in Table 3).

We repeat our previous simulation with a reduction of the parameter *V*_*de*_ by 10%, 20% and 30% with respect to its values listed in Table 1. Figure 5 shows the effect of reduction of *V*_*de*_ on the temporal variation of tau phosphorylation compared to the case where no PP2A reduction occurs as shown through solid line (0% curve). Although the qualitative nature of the curve remains the same but the reduced PP2A activity simulated via reduced value of the parameter *V*_*de*_ prevents the tau phosphorylation level (N) to restore to its homeostatic value (N_*h*_) even after a sufficiently long time. Figure 5 also shows that more the *V*_*de*_ deviation from its calibrated value (Table 1), the more is the residual tau phosphorylation level and so also the peak N value. The residual phosphorylation level of tau may be treated as an indicator of the irreversible phosphorylation of some tau protiens.

**Figure 5:**
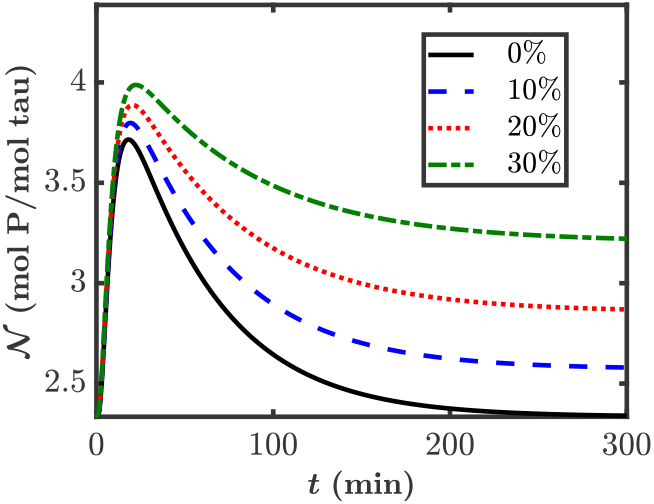
The effect of reducing the activity of PP2A on the evolution of tau phosphorylation levels. The loading case used here is CASE 1Int from Table 3. The reduction in the activity of PP2A phosphatase is simulated in our model via decrease in *V*_*de*_ value by 10%, 20% and 30% as from its calibrated value in Table 1.

We use our model along with spatial FEM simulations of brain injury over a 2D human brain after applying kinematic loads corresponding to the loading case 2 which results in only a mild TBI with slightly elevated peak tau hyperphosphorylation levels. However, this time, we predict the distribution of tau phosphorylation extent after three repeated TBIs. Each successive TBI is imparted via kinematic loading of the brain tissue after the effects of the previous injury have reached a steady state. Further, after each TBI, we progressively reduce the maximal tau dephosphorylation rate *V*_*de*_ by 20% uniformly throughout the brain. The peak tau phosphorylation levels thus obtained after each TBI incidence are plotted in Figure 6. Although the implementation of alteration of tau dephosphorylation due to injury is only rudimentary, we see that the peak tau phosphorylation levels progressively increase. Thus, our model predicts that any future TBIs will only lead to higher tau hyperphosphorylation levels, as is seen in neurodegenerative taupathies like CTE [8, 78]. Our model can, therefore, explain the increased vulnerability of axons to repeated mTBIs. Of note, we would like to clarify that due to absence of any data on mechano-sensitivity of tau dephosphorylation, we have assumed a uniform reduction in the parameter *V*_*de*_ after injury. In reality, the change being dependent on mechanical stress or the altered intra-axonal chemical signaling, is expected to be highly non-uniform, inducing higher non-uniformity in the tau phosphorylation distribution predicted by the model.

**Figure 6:**
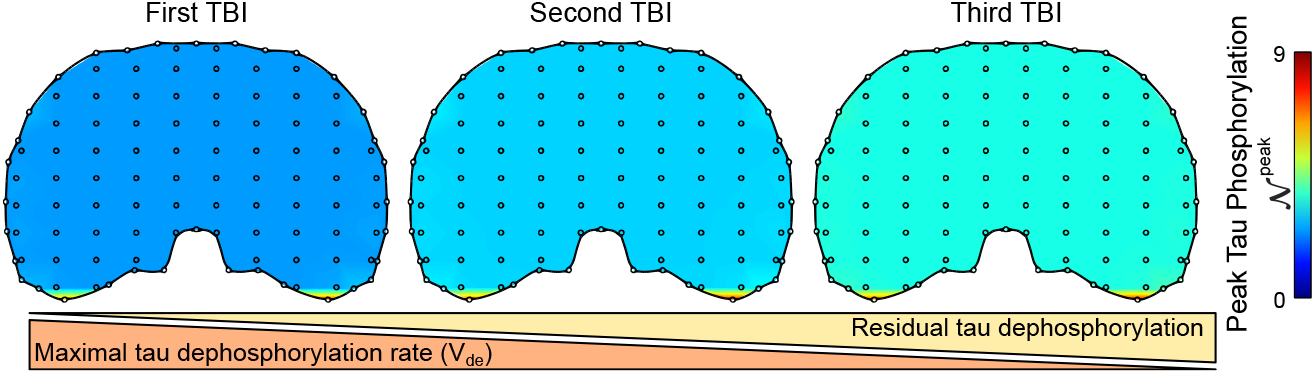
The effect of reducing the activity of PP2A on the evolution of tau phosphorylation levels. The loading case used here is CASE 1Int from Table 3. The reduction in the activity of PP2A phosphatase is simulated in our model via decrease in *V*_*de*_ value by 10%, 20% and 30% as from its calibrated value in Table 1.

## Discussion

After the occurrence of a TBI, the primary mechanical insult can result in generation of significant mechanical stresses in the brain tissue. However the physiological effects which characterize a TBI are typically an outcome of biochemical disturbances involved in a much delayed secondary insult phase. There have been extensive investigations into the mechanical nature of primary injuries and force transduction through the brain tissue and axons in predicting injury localizations. Yet, the role of mechanosensitive neurochemical disturbances coupled with the force mechanotransduction leading to chemo-mechanical axonal damage are largely unexplored. Here we present a time-dependent chemo-mechanical mathematical model to predict the extent of primary as well as secondary insult following a TBI.

While the secondary injury following TBIs can follow multiple pathways, leading to a highly variable pathophysiology, here we have focused on a prominent pathway initiated by injury-dependent calcium ion accumulation in the axons, leading to post-translational hyperphosphorylation of MT-binding tau proteins (Figure 1c). Following our previous work [29] where we had described the stress dependent calcium kinetics in a neuron experiencing an external mechanical impulse, we estimate the extent of calpain enzyme activation due to the intracellular *Ca*^2+^ accumulation. We next simulate the truncation of GSK-3*β* kinase by activated calpain, followed by the subsequent kinase-dependent phosporylation and phosphatase-dependent dephosphorylation of tau proteins. We provide a statistically averaged methodology to translate our results into a quantitative prediction of hyperphosphorylated tau. Lastly, we incorporate the spatial distribution of stresses in a human brain subjected to TBI-relevant kinematic loads, predicting the spatio-temporal tau-hyperphosphorylation based injury-maps following a TBI.

The delayed secondary insults of a TBI are initiated at an axonal level via a disturbance in the intracellular *Ca*^2+^ homeostasis. An excessive *Ca*^2+^ influx into the neuron via various plasma membrane (PM) transport channels and pumps, and an additional release of stored *Ca*^2+^ from the intracellular organelles like endoplasmic reticulum or the mitochondria result in an over-accumulation of *Ca*^2+^ inside the neurons (reviewed in [27, 79–81]). High levels of intracellular *Ca*^2+^ initiate a multitude of neurochemical reactions which result in production of free radicals, mitochondrial overload and disruption of essential cellular functions like glucose metabolism [80–83]. Intracellular *Ca*^2+^ also activates various degenerative proteases which act on readily available protein substrates throughout the neuron, compromising its structural and functional integrity, leading to fatal downstream effects such as activation of calpain.

Calpain is a family of cysteine proteases activated by *Ca*^2+^, of which calpain-I and calpain-II are ubiquitously present in human and animal tissues [45, 84, 85]. Calpain-I, which is more abundant in neurons, typically requires 3 − 50 *µ*M *Ca*^2+^ concentration for half-maximal activity and hence is also called *µ*-calpain. Calpain-II, also called m-calpain, on the other hand, is more expressed in glial cells and has a much higher half-maximal activity *Ca*^2+^ requirement (0.4 − 0.8 mM) [84]. Therefore, in this work, we focus on the proteolytic effects of *µ*-calpain. The *Ca*^2+^ requirement for calpain activation observed through experiments is however much higher than the physiological intracellular *Ca*^2+^ concentration, indicating a reduced in-vivo *Ca*^2+^ requirement, the exact reason for which is yet unresolved. It is hypothesized that in response to *Ca*^2+^ influx, calpain relocates to the cell periphery, where phospholipids present in the plasma membrane (PM) reduce the *Ca*^2+^ requirement for calpain activation [45, 84, 85]. It is also possible that limited activation of calpain at the locations of higher *Ca*^2+^ concentration is sufficient for downstream proteolysis of calpain substrates [45]. Notwithstanding the origin of calpain activation, intraneuronal calpain enzyme dependent proteolysis is widely implicated in secondary insult. Hence, we model the calpain activation as a single step reaction involving a cooperative binding of *Ca*^2+^ at approximately 6 active sites [42, 46, 47].

Within the axonal cytoskeleton, tau proteins can undergo various forms of post-translational modifications after a TBI, thereby losing their MT binding capabilities [8, 55, 59, 86]. Among these post-translational modifications, the phosphorylation performed by kinases and dephosphorylation by phosphatases are relevant to tau dysfunction. A delicate balance between tau phosphorylation and dephosphorylation is normally maintained so as to optimize the promotion of MT assembly. There are several kinases which phosphorylate tau in-vitro as well as in-vivo. These include glycogen synthase kinase-3*β* (GSK-3*β*), cyclin-dependent-like kinase 5 (CDK5), etc. Some kinases like protein kinase A (PKA) are known to prime the tau proteins for further phosphorylation by other kinase [34]. GSK-3*β* truncation in an AD affected brain correlates with tau hyperphosphorylation [40, 41, 59]. Similarly in a human TBI, the injury severity is found to correlate with increased activity of GSK-3*β* [54, 55]. Upon exposure to calcium, GSK-3*β* in human brain extracts is cleaved via a calpain mediation [35, 40, 56, 87]. Ample evidence is available in literature to implicate calpain in the GSK-3*β* truncation [35, 40, 56–59].

Reports suggest that GSK-3*β* in human brain extracts can be cleaved at N-terminal and/or at the C-terminal upon exposure to calcium [35, 40, 87]. Analogous to ΔN-GSK-3*β*, the truncation products ΔC-GSK-3*β*, and ΔN/ΔC-GSK-3*β* too show an increased activity and significantly contribute in tau phosphorylation. Currently our model (equations (5)) is solely based on ΔN-GSK-3*β* but, with the relevant kinetic data, can easily be extended to include the effect of ΔC-GSK-3*β*, and ΔN/ΔC-GSK-3*β* fragments.

There are several phosphatases which dephosphorylate tau, though protein phosphatase PP2A alone accounts for ∼ 70% of total phosphatase activity [36]. A calpain mediated disturbance in the signaling pathways causes a phosphorylation/dephosphorylation imbalance, by increasing the kinase activity while simultaneously decreasing the phosphatase activity, resulting in formation of hyperphosphorylated tau (P-Tau). Figure 1b shows a schematic representation of such an imbalance occurring due to a tauopathy, leading to P-tau formation. P-Tau in such a state is unable to bind to tubulin, thereby losing its ability to promote MT assembly. It has been observed that P-Tau further disrupts the MT assembly by sequestering normal tau from it [88]. Sequestered P-Tau nucleate the aggregation of more P-Tau into neurofibrillary tangles (NFT), as shown in Fig. 1c. The hyperphosphorylated tau oligomerize and aggregate, while the MT assembly disintigrates.

NFTs are observed commonly in multiple mild TBIs involving concussive or sub concussive impacts sustained over a period of years often observed among contact sport athletes and military veterans because of a high risk of purposeful, repetitive hits to the head [8, 78]. A similar tau protein dysfunction, via its hyperphosphorylation, is characteristically observed in a range of neurodegenerative disorders such as Alzheimers’ disease (AD), frontotemporal dementia, etc., together categorized as “tauopathies” [8, 39, 55, 89]. Repetitive mild TBI presents a major risk for development of CTE and other neurodegenerative tauopathies such as AD and dementia [8]. Tauopathies are characterized by abnormal hyperphosphorylation of tau protiens forming NFT aggregates. A human tau protein has upto 37 phosphorylation sites, though in a normal brain on an average only 2-3 moles of phosphates are present per mole of tau [36, 38, 60, 63]. Hyperphosphorylated tau (P-Tau) in an AD affected brain has 3-4 fold more phosphates per mole of tau [38, 62]. Thus, we believe that with appropriate adjustments, our kinetic model could also play a useful role in laying the foundation for a computational model studying tau hyperphosphorylation in such tauopathies.

## Conclusion

Following our previous work [30] where we had described the stress-dependent calcium kinetics in a neuron experiencing an external mechanical impulse, here we estimate the extent of calpain enzyme activation due to the intracellular *Ca*^2+^ accumulation. We also simulate the downstream proteolytic activity of calpain resulting in kinase activation followed by tau hyperphosphorylation, which leads to loss of MT-binding function of tau and subsequent MT assembly disruption. The predictions of our model match well with the key features of experimental and clinical observations:

- In the absence of any external load, a state of homeostasis is maintained, such that no calpain is activated, no kinase is truncated, and tau phosphorylation levels stay at their homeostatic value, 𝒩 _*h*_ = 2.33 mol P/mol total tau.
- The predicted in vivo activated calpain concentrations fall within the range of activated calpain used in vitro.
- The increase in GSK-3*β* activity towards tau due to calpain mediated truncation is seen to be qualitatively similar to the experimental reports.
- *Ca*^2+^ accumulation up to ∼ 8 times the homeostatic value will not directly result in increased tau phosphorylation. However, as more and more *Ca*^2+^ is accumulated, levels of tau phosphorylation become increasingly sensitive to it.
- Measurement of primary injury metrics such as maximum stress or strain localization does not always coincide with injury localization due to secondary insult.
- Reduced PP2A phosphatase activity after TBI results in unrecoverable tau phosphorylation levels, which makes successive TBI progressively more dangerous.
- A single mild TBI may not result in enough phosphorylation to observe any physiological symtomps. However, repeated incidences will result in an appreciable level of unrecoverable tau phosphorylation.

We believe our model provides a firm mathematical framework with a rich scope for further development. Either due to simplifying assumptions or due to lack of experimental data, the current model has a few limitations that could form the basis of any future work in this direction. Calpain mediated breakdown of other proteins such as spectrin could be incorporated in the model. The PM permeability is known to be altered due to spectrin breakdown. By carefully integrating an experimental evidence based functional dependence of membrane permeability on spectrin breakdown, the model can be enabled to capture a biphasic calcium accumulation leading to a sustained calpain mediated tau damage. Although we have incorporated the effect of reduction in the PP2A activity through parameter modification, an experimental investigation into the mechanism of PP2A inhibition after TBI is required to enhance the model, and predict the progressive vulnerability to successive injuries with a higher fidelity. Finally, to predict the extent of tau phosphorylation, we calculate the average number of occupied sites per tau proteins, considering all tau proteins whether phosphorylated or not. Establishing a relationship between the stochastic average and the number of tau proteins actually being phosphorylated will allow a better prediction of tau hyperphosphorylation. Lastly, coupling our model with a more detailed three-dimensional FE model of human brain would improve our predictive accuracy making the model practically useful for clinical applications.

## Author Contributions

AK, NVM and TKB conceived the study while NVM and TKB supervised it; AK performed simulation studies and analyses; AK, NVM and TKB wrote the manuscript.

## Footnotes

1 kiloDaltons, denoting mass

