## Supplementary Material for "Chemo-Mechanical Regulation of Tau Phosphorylation Following Traumatic Brain Injuries"

### 1 An Interpretation of Tau Phosphorylation Level ($\mathcal{N}$) for Future Scope

Our model predicts the level of phosphorylation $\mathcal{N}$ defined as the average number of phosphorylated sites per tau protein upon the action of external mechanical load incurred during incidences like TBIs. Physiologically the effects of tau phosphorylation are seen in the form of reduced affinity of hyperphosphorylated tau towards the MT bundles, thereby disrupting the MT assembly, leading to formation of axonal varicosities [8, 9, 1]. The intensity of physiologically observed secondary injuries might be quantified in terms of the number of hyperphosphorylated tau proteins. As pointed earlier, the present work sidesteps the stochastic nature of the tau phosphorylation process and offers an average prediction. A means to rectify this limitation is through an assumed probability distribution of the phosphorylation level amongst all the tau proteins, such that the mean of the probability distribution corresponds to $\mathcal{N}$, the average occupancy per tau protein as predicted by our model at any time. Given a probability density function PDF ($\mathcal{N}_x$) over the randomly distributed phosphorylation levels $\mathcal{N}_x$, we can calculate the fraction of tau proteins hyperphosphorylated by introducing a threshold value of $\mathcal{N}$, such that tau proteins with phosphorylation levels above this threshold value, say $\mathcal{N}_{th}$, are considered to be hyperphosphorylated. Amongst the many possible forms for the probability distribution function (PDF), the one that is selected should be in sync with the problem at hand. As a first approximation, we choose the widely applied PDF *log-normal distribution* which is skewed, usually used in situations involving low mean values, large variances, non-negative value of the random variable [6]. The estimation of the variance of the PDF may be feasible using experimental measurement of the phosphorylation levels of in-vitro brain tissue specimens undergoing a simulated injury.

For a log-normal PDF,

$$\text{PDF}(\mathcal{N}_x) = \frac{1}{\mathcal{N}_x v\sqrt{2\pi}} e^{-\frac{(log(\mathcal{N}_x)-m)^2}{2\times v^2}} \tag{1}$$

the fraction of hyperphosphorylated tau proteins are given as,

$$\frac{[\text{P-tau}]}{[\text{tau}]_0} = \int_{\mathcal{N}_{th}}^{\infty} \left\{ \frac{1}{\mathcal{N}_x v\sqrt{2\pi}} e^{-\frac{(log(\mathcal{N}_x)-m)^2}{2\times v^2}} \right\} d\mathcal{N}_x \tag{2}$$

where, $m$ and $v$ are parameters of the probability distribution governing its median and skewness respectively. We consider a log normal PDF with mean at $\mathcal{N} = 3$ and assume for the sake of calculations $v = \log(1.5) = 0.4$, a value typically observed in a variety of medical and epidemiological observations [6]. Further we decide that hyperphosphorylation of tau occurs over a threshold level $\mathcal{N}_{th} = 6$ mol P/ mol tau, based on the lower limit of phosphorylation levels observed in AD P-tau [5, 2]. Through equation (2), we predict that when the phosphorylation levels reach peak value $\sim 7\%$ tau would be hyperphosphorylated. The reduction in MT assembly stability post tau removal due to hyperphosphorylation has been previously studied, though without a motivation for the number of tau proteins being removed [7]. A mathematical framework, motivated by our speculative discussions in this section, permits the prediction of the number of hyperphosphorylated tau proteins, thus bridging an important gap. It goes without saying that experimentally supported parameter values and PDF would yield a better prediction of the fraction of hyperphosphorylated tau.

### 2 Additional figures and tables

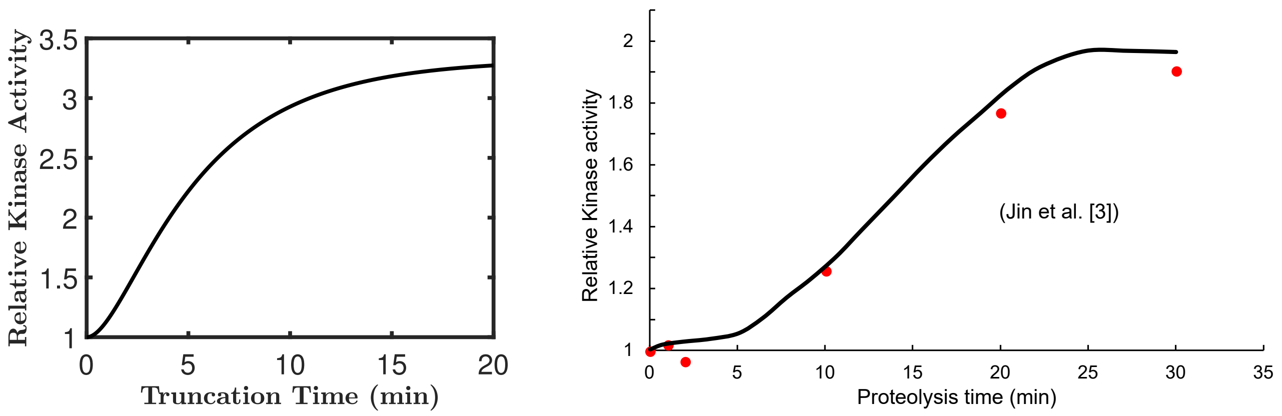


**Figure S1:** (a)The simulated relative increase in the GSK-3$\beta$ phosphorylation activity towards tau based on Equations (6) and (8) after the truncation of in-silico incubation with 1nM calpain-I at various times. (b) A qualitatively similar increase in kinase activity observed by Jin et al. [3] due to truncation of GSK-3$\beta$ with calpain for the specified proteolysis time in-vitro.

**Table S1:** Value of Calcium Kinetics Parameters [4]

| Parameter | Definition | Value |
|---|---|---|
| $c_i^*$ | Homeostatic value of intracellular concentration | $1.00 \times 10^{-4}$ mM |
| $c_e^*$ | Homeostatic value of extracellular concentration | 1.00 mM |
| $c_{\text{er}}$ | $Ca^{2+}$ concentration inside the ER | 0.10 mM |
| $K_{\text{pm}}$ | Porosity of the PM | $2.94 \times 10^{-6}$ s$^{-1}$ |
| $K_{\text{er}}$ | Porosity of the ER membrane | $3.17 \times 10^{-5}$ s$^{-1}$ |
| $k_{\text{pm}}$ | Activation constant of the PM | $2.00 \times 10^{-4}$ mM |
| $k_{\text{er}}$ | Activation constant of the ER membrane | $5.00 \times 10^{-4}$ mM |
| $n_{\text{pm}}$ | Hill Coefficient of the PM | 2.00 |
| $n_{\text{er}}$ | Hill Coefficient of the ER membrane | 1.00 |
| $\chi_{\text{pm}}$ | Stress dependence parameter for $V_{\text{pm}}$ | $2.00 \times 10^3$ |
| $\chi_{\text{er}}$ | Stress dependence parameter for $V_{\text{pm}}$ | $4.00 \times 10^3$ |
| $\kappa$ | Stress dependence parameter 2 | $4.50 \times 10^{-5}$ Pa$^{-1}$ |
| $\alpha$ | Impulse contribution parameter | $7.5 \times 10^{-3}$ s$^{-1}$ |
